# Human keratinocyte (HaCaT) stimulation and healing effect of the methanol fraction from the decoction from leaf from *Sideroxylon obtusifolium* (Roem. & Schult.) T.D. Penn on experimental burn wound model

**DOI:** 10.1101/2020.03.29.013839

**Authors:** Tamiris de Fátima Goebel de Souza, Taiana Magalhães Pierdoná, Fernanda Soares Macedo, Pedro Everson Alexandre de Aquino, Gisele de Fátima Pinheiro Rangel, Rebeca Silva Duarte, Glauce Socorro de Barros Viana, Ana Paula Negreiros Nunes Alves, Raquel Carvalho Montenegro, Diego Veras Wilke, Edilberto Rocha Silveira, Nylane Maria Nunes de Alencar

## Abstract

The larger number of plants, with therapeutic potential, popularly used in Northeastern Brazil is due to their easy access and the great Brazilian biodiversity. Previously, was demonstrated that the methanol fraction from *Sideroxylon obtusifolium* (MFSOL) promoted an anti-inflammatory and healing activity in excisional wounds. Thus, this work aimed to investigate the healing effects of MFSOL on human keratinocytes cells (HaCaT) and experimental burn model injuries. HaCaT cells were used to investigate migration and proliferation of cell rates. Female Swiss mice were subjected to second-degree superficial burn protocol and divided into four treatment groups: Vehicle (cream-base), 1.0% Silver Sulfadiazine (Sulfa), and 0.5% or 1.0% MFSOL cream (CrMFSOL). Samples were collected for quantification of the inflammatory mediators and histological analyses after 3, 7 and 14 days on evaluation. As result, MFSOL (50 μg/ml) stimulated HaCaT cells by increasing proliferation and migration rates. Moreover, CrMFSOL 0.5% attenuated myeloperoxidase (MPO) activity and also stimulated the release of IL-1β and IL-10, after 3 days with treatment. CrMFSOL 0.5% enhanced wound contraction, promoted tissue remodeling improvement and highest collagen production after 7 days, and VEGF release after 14 days. Therefore, MFSOL evidenced the stimulation of human keratinocyte (HaCaT) cells and improvements on wound healing via inflammatory modulation on burn injuries.

## 1. INTRODUCTION

The reconstruction of the epidermal barrier rupture through re-epithelialization is essential to reduce a wound infection risk and recover the normal skin function (Rousselle, Braye, & Dayan, 2018). The burn healing process is affected by interaction among cells and molecules, including growth factors, cytokines, and chemokines. Initially, the inflammatory response occurs to prevent the invasion of pathogens and the activation of fundamental signals. It is also followed by the proliferative phase, when fibroblasts and keratinocytes will restore the tissue damage through collagen synthesis, epithelialization, angiogenesis, and production of extracellular matrix (Horst, Chouhan, Moiemen, & Grover, 2018; Markeson et al., 2015).

The use of medicinal plants in Brazil is still common because of both high costs of traditional medicines and easy access to medicinal plants which are sold on flea markets of large and small cities (Macedo et al., 2018). *Sideroxylon obtusifolium* (Roem. & Schult.) T.D. Penn is a native tree in Central and South America. It is characteristic of the Brazilian semiarid region (Caatinga), where is known as “quixaba”, due to its round blackberries when ripened. This medicinal plant species is widely known and used in folk medicine in the cities of Northeast of Brazil, where dry parts of quixaba, leaves or trunk bark, are sold in local markets due to their anti-inflammatory and healing properties (Oliveira et al., 2012).

The risk of extinction of the species, by the predatory extractive procedures of its bark, directed the attention of scientific studies to other plant parts like the leaves. Two studies with the hydroalcoholic fractions obtained from leaves of *S. obtusifolium* showed antifungal activity against *Candida albicans* (Pereira et al., 2016; Sampaio et al., 2017). Other study demonstrated antinociceptive and anti-inflammatory properties of L-proline, *N*-methyl-(2S, 4R)-*trans*-4-hydroxy-L-proline (NMP), presents in major proportion in the methanol fraction of leaves of *S. obtusifolium* (MFSOL) (Aquino et al., 2017). Also, NMP showed wound healing activity on excisional wounds, which was related to its anti-inflammatory and antioxidant actions (Aquino et al., 2019). However, there is no knowledge of whether methanol fraction MFSOL is the healing process in wounds caused by burns. Then, this study aimed to investigate the effect of MFSOL on human keratinocytes (HaCaT) *in vitro* and on surface burn model in mice.

## 2. EXPERIMENTAL

### 2.1 Plant material

Leaves of *S. obtusifolium* (Sapotaceae) were collected from specimens growing in Mauriti city, state of Ceará, Brazil. A voucher specimen (sample no. 10.648) was taxonomically validated by Dr. Maria Arlene Pessoa da Silva, at the “Herbario Caririense Dárdano de Andrade Lima”, in Regional University of Cariri, Ceará, Brazil.

The methanol fraction from *S. obtusifolium* leaves (MFSOL) was obtained according to the procedure previously described, to obtain a yellowish amorphous powder (Aquino et al., 2017). The MFSOL was dissolved in sterile water and treated to cell cultures at final concentrations of 6.25 –100 μg/ml.

### 2.2 Cell Culture

Immortalized human keratinocyte cells (HaCaT, CLS Cell Lines Service) were provided by Dr. Raquel Carvalho Montenegro, Pharmacogenetics Laboratory of the Drug Research and Development Center – NPDM, Federal University of Ceará. Cells were grown in Dulbecco’s modified Eagle’s medium (DMEM, Gibco®), supplemented with 1.0% antibiotics (100 U/ml penicillin and 100 μg/ml streptomycin, Gibco®) and 10% fetal bovine serum (FBS, Gibco®), at 37°C in 5% CO_2_ atmosphere in a humidified incubator. The cells were maintained in exponential growth through periodic maintenance.

### 2.3 Cytotoxicity assays

The principle of MTT assay is based on the mitochondrial cell viability to reduce, by the succinate-tetrazole reductase enzyme system, the yellow tetrazolium salt (MTT) to a salt-formazan, which is purple in color (Mosmann, 1983). After 24, 48 and 72 h of incubation with MFSOL, cells received 20 μL of 3-(4,5-dimethylthiazolyl)-2,5-diphenyltetrazolium bromide (MTT) (5 mg/ml) and were incubated at 37 °C for 3 h. Then, the medium was removed and 150 μL of dimethyl sulphoxide (DMSO) was added and the plate was homogenized for 5 min. The absorbance was determined by a microplate reader (Elisa Asys Expert Plus) at a wavelength of 540 nm.

The sulphorhodamine B (SRB) assay was used to evaluate cell viability, based on the measurement of cellular protein content. HaCaT cells (2 × 10^4^ cells/ml) were plated in 96 multiwell plates and after 24h treated with MFSOL (6.25 – 100 μg/ml). After the 24, 48 and 72h of treatment, the cells were fixed with 10% trichloroacetic acid and incubated with SRB solution (0.4%) for 30 min. The dye excess was removed by washing repeatedly with 1% acetic acid. The protein-bound dye was dissolved in 10 mM Tris-base solution and the optical density was determined at 570 nm using a Microplate Autoreader (Multiskan FC - Thermo Scientific) (Houghton et al., 2007). The results were expressed as a percentage of cell viability.

### 2.4 Scratch wound healing assay

The scratch wound healing assay was performed to determine the effects of MFSOL on the proliferation and migration of human keratinocytes. These cells were dispersed in DMEM with 10% FBS (5 × 10^4^ cells/ml) and incubated with 5% CO_2_ at 37 °C in a 24 well plate. When the cells formed a confluent monolayer, they were scratched using a tip vertically with a 200 μl of each well. Cellular debris were removed and washed with PBS. Pictures (200x) were taken after 24, 48 and 72 h of incubation with MFSOL (25, 50 and 100 μg/ml) in 1ml of fresh medium (Räsänen & Vaheri, 2010). To evaluate only the migration, an antimitotic agent, mitomycin C (10 μg/ml; Sigma®) was added for 1 h, following the scratch protocol. Immediately, the cells were treated with MFSOL (25 and 50 μg/ml) and, after photomicrographs, the migration was evaluated at the initial time, and after 24 and 48 h of incubation (D. Choi, Piao, Wu, & Cho, 2013). The open area of the scratch was measured using TSCRATCH® software in each analysis period in the same initial site, the percentage of the open area was measured using the formula: Open area (%) = Open area time *X* ÷ Open area initial time ×100.

### 2.5 Animals

Female Swiss mice (8 weeks old) were kept in a light/dark 12h cycle, temperature (23 ± 2°C) and humidity (55 ± 10%) at Central Animal House of the Federal University of Ceará (UFC). The animals received food (Nuvilab, Quimtia®, Brazil) and water *ad libitum*. The experimental protocols were performed according to the ethical standards established in the Ethical Principles on Animal Experimentation adopted by the Brazilian National Council for the Control of Animal Experimentation (CONCEA), analyzed and approved (n° 8862290518) by the Ethics Committee on the Use of Animals of Federal University of Ceará.

### 2.6 Superficial burn model and treatment

Firstly, the animals were anesthetized with xylazine hydrochloride (10 mg/kg, i.p.) and ketamine hydrochloride (100 mg/kg, i.p.). After anesthesia, the dorsal surface skin (4 cm × 2 cm) was shaved followed by asepsis with 1.0% iodopovidone followed by 70% ethanol. A second-degree superficial burn was induced by direct contact of a heated square stainless steel plate (1 cm^2^) and regulated at 100 °C for 6 seconds (Lima-Junior et al., 2017). The animals were divided into 4 groups: 1) control vehicle with cream base Lanette (anionic cream); 2) CrMFSOL 0.5% and 3) CrMFSOL 1.0%, which fraction was included in the base dermatological cream in two concentrations, respectively; and 4) 1% silver sulfadiazine with cream base Lanette as the positive control. Immediately after the injury, the treatments were administered once per day for 14 days. After induction of the lesion, the animals received saline solution 0.9% (s.c.) for fluid replacement and were kept in a warm environment and under observation until complete recovery.

### 2.7 Burn wound contraction

The lesions were measured and photographed on 3, 5, 7, 9, 12 and 14 days after induction of the superficial burn. A digital caliper was used to measure the horizontal (h), and vertical (v) measurements of the burn (mm), and then the area of the lesion was calculated by multiplying the two measurements, A = h × v (mm^2^). The rate of contraction of the lesion was calculated by the formula: Contraction (%) = (initial area – evaluation area) / (initial area) × 100. After macroscopic wound healing examination (n=10/group), all mice were euthanized using anesthetic overdose (xylazine and ketamine) and tissues were collected immediately and kept at −80° C for biochemical analyses.

### 2.8 Histological and collagen fibers analysis

The burn injuries were collected after 7 days of burn injury to histological study. Samples were removed and fixed in 10% formol (pH 7.4) for 24 h (n=6/group). Tissues were submitted to dehydration and impregnated in paraffin. Sections of those fragments were cut to 4 μm in thickness and stained with hematoxylin and eosin. Histopathological changes were evaluated with optical microscopy (Olympus BX 51, Japan). The histopathological parameters were determined and scored from 0 to 4,where zero score corresponding: absence of ulcer (AU), remodeled connective tissue (RCT); score 1: AU, fibrosis (F), slight chronic inflammation (CI); score 2: presence of ulcer (PU), F, moderate CI; score 3: PU, chronic inflammation process (granulation tissue) and score 4: PU, acute process (dilated vessels, mixed inflammatory infiltrate with neutrophils) (Adapted from Cavalcante et al., 2011). Stained with picrosirius red (PSR) were performed to measure the collagen fibers in the connective tissue of burn wounds. The Color Deconvolution (RGB) plugin of the Image J® software was used to measure the percentage of collagen area represented by the red image produced concerning the total area of the image (Alves et al., 2015).

### 2.9 Myeloperoxidase activity assay

The myeloperoxidase (MPO) activity was performed in burns biopsies after 3 days of burn injury (Adapted from Faunce et al., 1999). The samples were homogenized using a polytron tissue extractor in phosphate buffer ph 7.4, centrifugated ate 10.000 rpm for 30 min and the pellet obtained here used to determine myeloperoxidase activity by oxidative reaction with 3,3’,5,5’-tetramethylbenzidine (TMB; 1,6 mM, Sigma®) and oxygen peroxide (H_2_O_2_; 0,5 mM). The number of neutrophils was quantified from a standard neutrophil curve (1× 10^5^ neutrophils/ well). The absorbance of the samples (n=6/group) was quantified in a spectrophotometer at the wavelength of 450 nm, and the results were expressed as the number of neutrophils/mg of tissue (cell/ mg of tissue).

### 2.10 Mediators of inflammation and vascular endothelial growth factor measurement

The biopsies of the burns were performed after 3 days of burn injury to quantify the release of inflammatory mediators TNF-α, IL-1β, IL-10 and after 14 days of growth factor VEGF (Fujimi et al., 2009). Tissues were triturated and homogenized 10% (mg tissue/μL) at 4 °C in PBS solution (pH 7.4), and the residues removed after centrifugation at 5000 rpm for 5 min. The protocol was performed according to the manufacturer’s recommendation to the conventional sandwich technique (R&D Systems®). After the ELISA protocol, the absorbance of the samples was quantified in a spectrophotometer at the wavelength of 450 nm. The results were expressed as cytokines/ml of homogenate, and the concentration of the samples was obtained from a serial dilution of recombinant cytokine standard (n=6/group).

### 2.11 Statistical analysis

Results are presented as the mean ± standard deviation (SD) for each experimental group. Statistical comparisons of the data were performed by one-way ANOVA followed by Tukey post-test using GraphPad Prism software version 5.0. Differences between groups were considered significant when *p* ≤ 0.05.

## 3. RESULTS

### 3.1 MFSOL enhanced the cellular viability of human keratinocyte HaCaT

The proliferative assay evidenced that MFSOL did not show cytotoxicity in human keratinocytes, at any of the incubation periods. MFSOL was able to increase mitochondrial viability (by MTT assay) after 24, 48 and 72 h at concentrations of 12.5 - 50 μg/ml (p<0.001). The concentration 50 μg/ml of MFSOL stimulated an increase in the viability of 42.8% (after 24 h), 47.5% (after 48 h) and 46.8% (after 72 h) compared to the Control group (p <0.001) (Fig. 1A, 1B and 1C, respectively). At the highest concentration (100 μg/ml) it showed an increase of 43.6% (p<0.001) only after 24 h. Following, the SRB assay revealed an increase of viable cells, but less expressive than MTT assay. After 24 h and 48 h, the concentration 100 μg/ml of MFSOL increased 14.5% and 21% cell viability of HaCaT cells (p<0.001), what also was observed with 50 μg/ml (8.2% and 11.21%; p<0.001), respectively. These effects were less pronounced after 72 h when at 25 and 50 μg/ml of MFSOL showed an increase of 7.1% and 7.4% (p<0.001) of viability.

**Figure 1.**
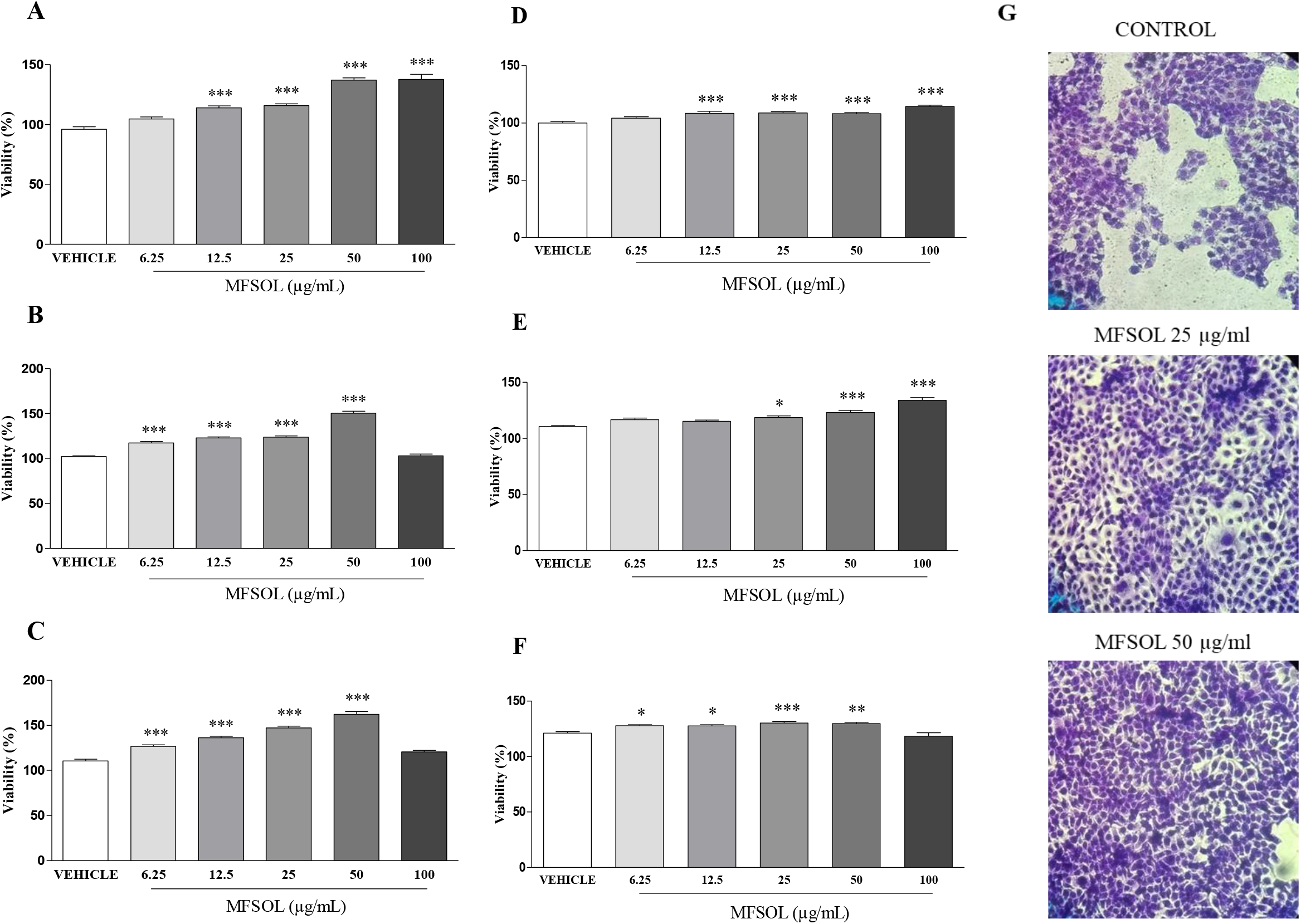
Effect of MFSOL on cell viability of human keratinocytes (HaCaT). An increase of cell viability of keratinocytes was produced by MFSOL (6.25 - 100 μg/mL) after 24h, 48h, and 72h incubation, by MTT (A, B, and C) and SRB (D, E, and F) methods, respectively. To illustrate the HaCaT density, photographs were taken at 200x magnification after 48 hours of treatment with 25 and 50 μg/ml of MFSOL (G). Values were represented as the mean ± SE from at least three independent experiments. \**p*<0.05, \*\**p* <0.01 and \*\*\**p* <0.001, compared with Vehicle group.

### 3.2 MFSOL stimulates cell proliferation and migration in the HaCaT scratch assay in keratinocytes

To evaluate the effect of MFSOL on cell proliferation and migration of HaCaT, the *in vitro* healing assay known as Scratch was used. Through the MTT and SRB assays, concentrations of 25, 50 and 100 μg/ml were selected to perform Scratch. After 24 h, the percentage of the open area observed after treatment with 25 μg/ml (p<0.05) and 50 μg/ml (p<0.001) of MFSOL was reduced by 14.5% and 34%, respectively, compared to the Control (Fig. 2A). This effect was also observed after 48 h after treatment with 12.5% (22%; p<0.05), 25 μg/ml (28%; p<0.001) and 50 μg/ml (38%; p<0.001) of MFSOL, respectively, when compared to Control (Fig. 2B). Finally, after 72 h of the scratch assay, the open area was reduced by 25 μg/ml (42.1%; p<0.05) and 50 μg/ml (72.6%) of MFSOL (p<0.001), compared to Control (Fig. 2 C). The effect of MFSOL on HaCaT was photographed and visualized through rapid panotype staining. It was observed by the higher cell density in 25 μg/ml and 50 μg/ml of MFSOL treatment in comparison to the Control (Fig. 2D). Considering the stimulatory effect of MFSOL on cell proliferation and migration, mitomycin C was used to inhibit cell duplication and thus enable the isolated evaluation of the effect of MFSOL on cell migration. At the concentration of 50 μg/ml was efficient in stimulating the migration of the cells to the center of the open area by 26% after 24 h (p<0.05) and 67% after 48 h (p<0.01) of incubation, compared to Control (Fig. 2F-G).

**Figure 2.**
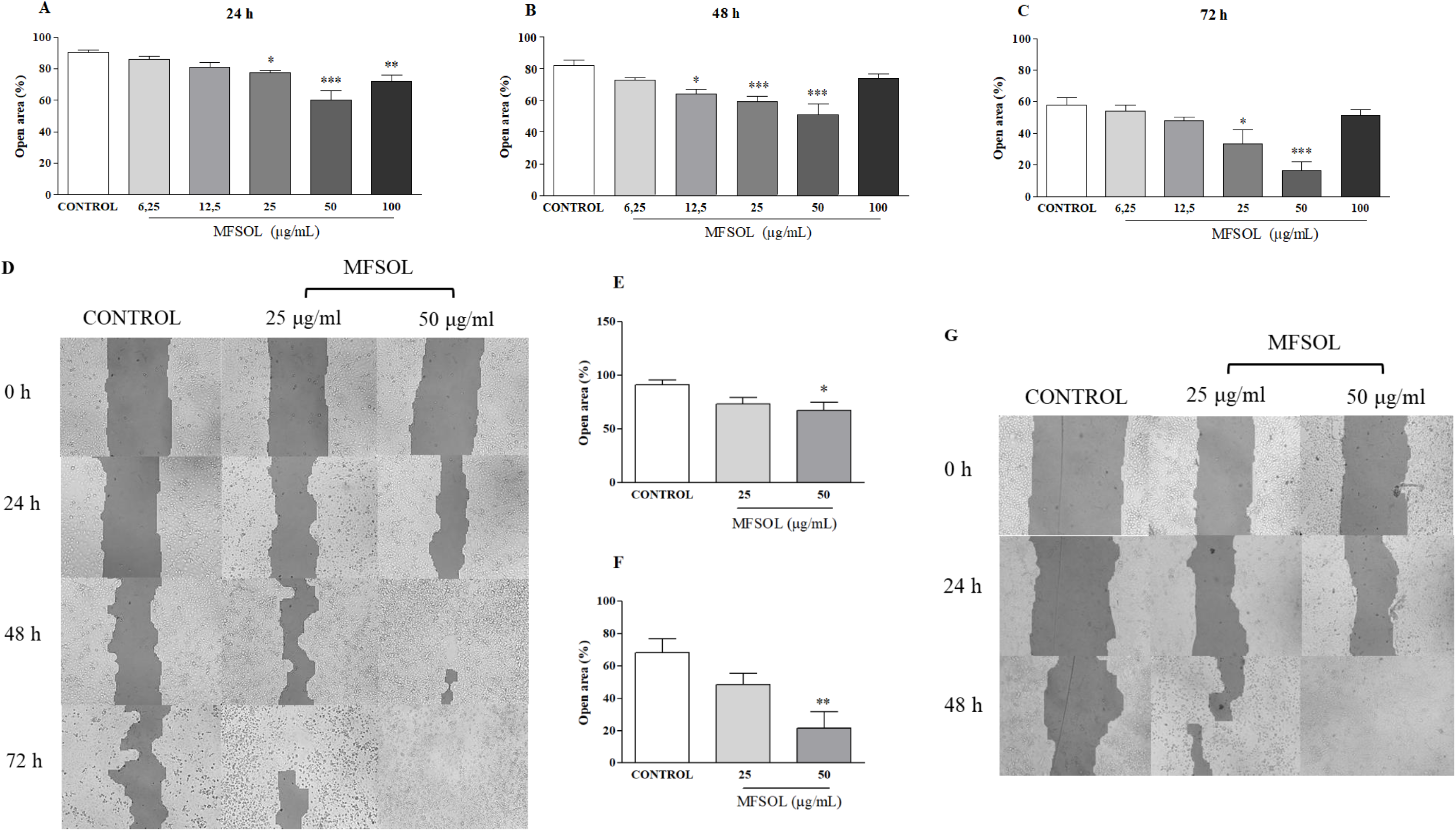
MFSOL stimulated proliferation and cell migration rates of HaCaT in the Scratch assay. A scratch was produced in a monolayer of HaCaT cells and photographs were taken, before (0h) and after 24 h (A), 48 h (B) and 72 h (C) of treatment with MFSOL, with a 200 × magnification (D). As cells double and migrate, space is filled and the open area is reduced. A pre-treatment with mitomycin C (10 μg/ml) was also performed one hour before Scratch to inhibit cell proliferation, allowing only evaluation of cell migration after 24 h (E) and 48h (F). The wells were photographed (200x) after 24 h and 48 h (G). Photomicrographs are produced by the open area analysis (%) through the TSCRATCH® program. The results were expressed as mean ± standard error of the mean. One-way ANOVA followed by the Tukey test was used for statistical analysis of the data. \**p*<0.05, \*\**p* <0.01 and \*\*\**p* <0.001 represent a significant difference compared to the Control group (water).

### 3.3 CrMFSOL reduces the area of superficial burns

In the superficial burn model, the contraction of lesions was measured for 14 days to monitor the temporal evolution of the area of superficial burns. The contraction rate of the Sham and Vehicle groups was similar in all the periods studied (Figure 3A-F). After 3 days, the treatment with CrMFSOL 1.0% showed an increase by 123.2% (p<0.05) in the contraction of lesions, while, the Sulfa group (p<0.05) also presented similar effect (110.7%), compared to the Vehicle group (Fig. 3A). Following after 5 days (Fig. 3B) and 7 days (Fig. 3C), CrMFSOL 0.5% (p<0.01; p<0.05), CrMFSOL 1.0% (p<0.05; p<0.01) and Sulfa (p<0.01; p<0.01) groups showed an increase of lesions contraction, respectively, compared to the Vehicle group (Fig. 3C). At 9 days, only CrMFSOL 0.5% group increased (26%) the lesions contraction (p<0,05) when compared to Vehicle (Fig. 3D). Near the closure of lesion, after 12 and 14 days of analysis, all the groups presented no differences between them (Fig. 3E and 3F). The photographs represent the evolution during the cicatricial process of the studied times 3rd (A), 5th (B), 7th (C), 9th (D), 12th (E) and 14th (F) day (Fig. 3G), respectively.

**Figure 3.**
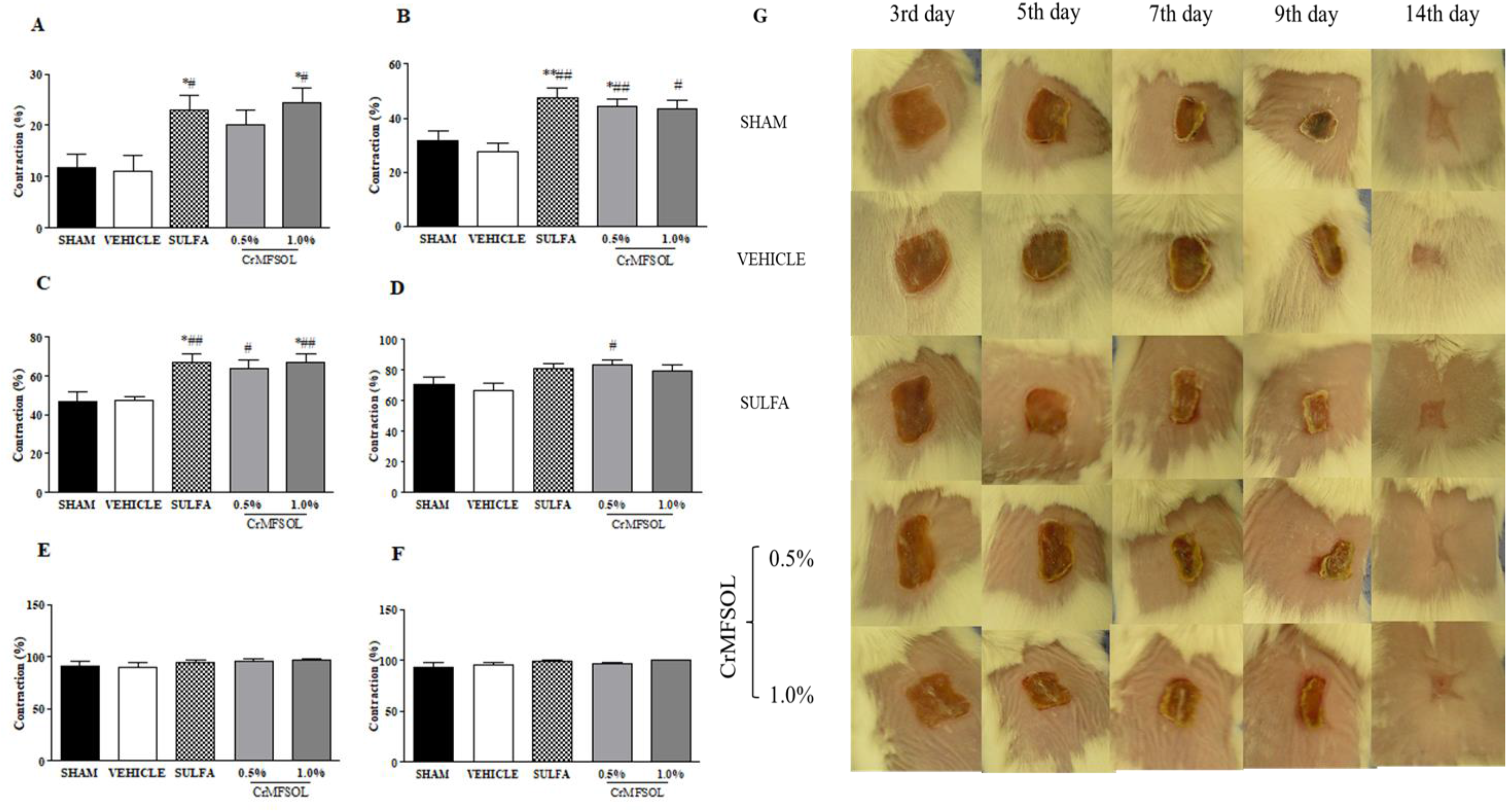
Effect of MFSOL on the contraction of induced burns in mice. The percentage of contraction of the burns was expressed after the 3, 5, 7, 9, 12 and 14 days (A, B, C, D, E, and F, respectively). In the initial days, the wound healing of the groups treated with 0.5% and 1.0% CrMFSOL was faster. The animals were treated with a single daily application of dermatological creams for 14 days, the CrMFSOL group corresponds to the 0.5% and 1.0% of MFSOL, the Sulfa group to the 1% silver sulfadiazine cream and Vehicle to the base vehicle cream without any addition of active substance. The superficial burns were photographed for macroscopic monitoring on days 3, 5, 7, 9, 12 and 14 after the burn was produced (G). One animal per group was chosen to represent the group according to the results obtained by the contraction of the lesion. One-way ANOVA followed by the Tukey test was used for statistical analysis of the data. \**p* <0.05 and \*\**p* <0.01 represents a significant difference compared to the Sham group and ^#^*p* <0.05 and ^##^*p* <0.01 for the Vehicle group, (n = 10 animals / group).

### 3.4 CrMFSOL promotes tissue remodeling improvement, reepithelialization and collagen deposition on superficial burns

Histopathological analysis of samples of superficial burns in mice showed that after seven days the Sham and Vehicle groups had ulcer, granulation tissue and presence of acute inflammatory infiltrate (Fig. 4A and 4B). On the other hand, Sulfa, CrMFSOL 0.5%, and 1.0% (p<0.05) groups stimulated the appearance of a thin epithelium covering the ulcer, with the presence of a corneal layer protecting the newly formed epithelium. There was still the transition from acute to chronic inflammatory infiltrate and the development of a remodeling connective tissue (Fig. 4C, 4D and 4E, respectively). To determine the total collagen deposition in the tissue, the staining of picrosirius red was used after 7 days of treatment. The Vehicle and Sham groups had a similar percentage of total collagen. While Sulfa, CrMFSOL 0.5% and 1.0% groups were increased 36%, 30.7% and 48.1% (p<0.05), respectively, the production of collagen in comparison to the Vehicle group (Figure 4F).

**Figure 4.**
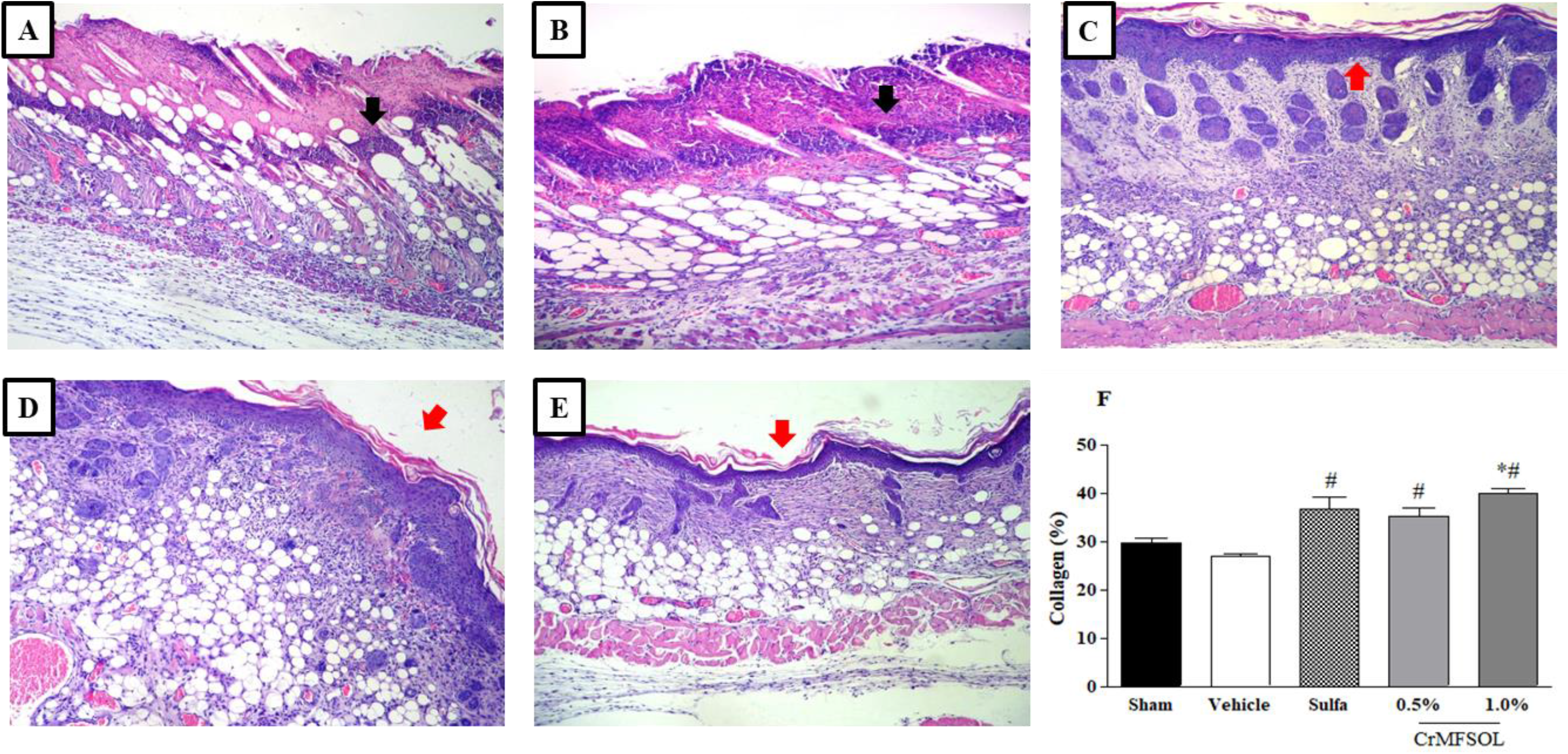
MFSOL promoted tissue remodeling improvement, reepithelialization and stimulated collagen production on superficial burns after 7 days of treatment. Photomicrographs of histological slides (Hematoxylin Eosin stain) from burn lesions on the 7th day after surgery (100x) were represented by Sham (A); Vehicle (B); Silver Sulfadiazine (C), 0.5% CrDFSO-M (D) and 1.0% CrDFSO-M (E) groups (n = 6 animals/group). Black arrows indicate the presence of acute inflammatory infiltrate in groups A and B, while red arrows indicate the presence of atrophic epithelium, presence of a corneal layer and absence of ulcer in groups C, D and E. The lesions were stained by the picrosirius red method and photographed in six fields (200x). The percentage of total collagen present on the 7th day was determined by Image J® software (F). The results were expressed as the mean of the six areas of each wound per group. One-way ANOVA followed by the Tukey test was used for statistical analysis of the data. \**p* <0.05 corresponds to the difference in relation to the Sham group and ^#^*p* <0.05 in the Vehicle group (n = 6 animals/group/day).

### 3.6 CrMFSOL on cytokine inflammatory status in superficial burns

The treatment with 0.5% CrMFSOL group reduced 32% (p<0.05) the activity of the myeloperoxidase (MPO) when compared to the Vehicle group after 3 days of burn-induced injury (Fig. 5A). Moreover, the levels of TNF-α were not altered by CrMFSOL 0.5% or 1.0% (Fig. 5B). However, CrMFSOL 0.5% (p<0.001) and 1.0% (p<0.01) CrMFSOL increased IL-1β release (112.5% and 94.4%, respectively) compared to the Vehicle group, what was also observed with the Sulfa group treatment (p<0.001) (Fig. 5C). Additionally, the release of IL-10, an anti-inflammatory cytokine, was higher in all treated groups compared with the Vehicle group. CrMFSOL 0.5% treatment showed a considerable increase of 153.7% (p<0.001) and 59.8% (p<0.01) when compared to the Vehicle and Sulfa groups, respectively (Fig. 5D).

**Figure 5.**
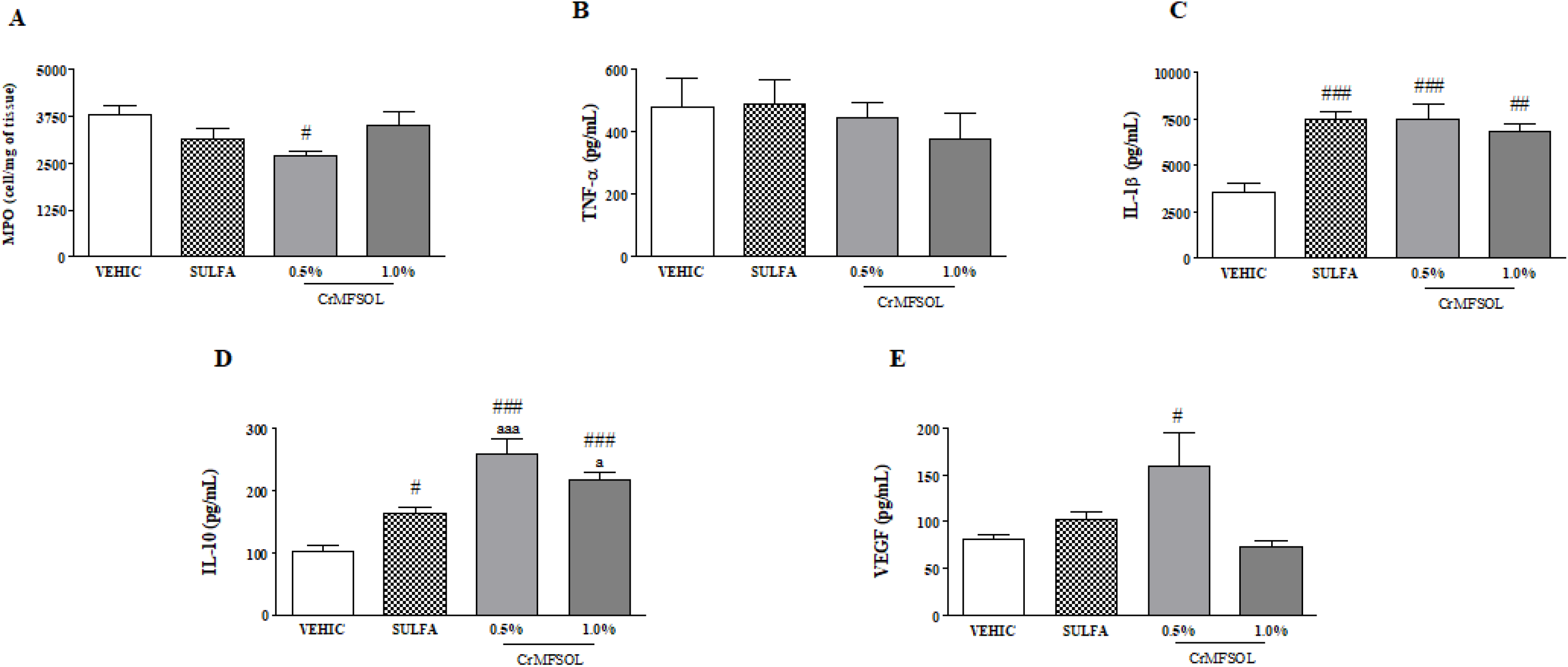
The treatment with MFSOL reduced myeloperoxidase activity (MPO) and stimulated release of IL-1β, IL-10 after 3 days and VEGF after 14 days. The levels of TNF-α were not reduced significantly. The results of MPO (A) were expressed as mean ± standard error of the mean of cell/mg of tissue, while TNF-α (B), IL-1β (C), IL-10 (D) and VEGF (E) were expressed as the mean ± standard error of the mean of cytokine/ml from supernatant/mg tissue. One-way ANOVA followed by the Tukey test was used for statistical analysis of the data. ^#^*p* <0.05, ^##^*p* <0.01 and ^###^*p* <0.001 represents a significant difference in comparison to Vehicle group. ^a^*p* <0.05 and ^aaa^*p* <0.001 in relation to the Sulfa group (n = 6 animals/group/day).

### 3.7 CrMFSOL stimulated release of VEGF on superficial burns

To evaluate the angiogenesis process, the VEGF level was measured after 14 days of induced superficial burn in animals. The treatment with CrMFSOL 0.5% was able to increase VEGF release by 95.6% (p<0.05) compared to the Vehicle group (Fig. 5E).

## 4. DISCUSSION

This work reports, for the first time, the healing activity on burn wounds of methanol fraction from *Sideroxylon obtusifolium* (MFSOL). Regarding the traditional use of stem bark, this work sheds light on a sustainable exploration of *S. obtusifolium* by shifting to, the use of its leaves. MFSOL was not cytotoxic and promoted an increase of cellular viability of human keratinocytes (HaCaT), suggesting an effect on the proliferation of HaCaT. Similarly, the hydroalcoholic extract of the trunk bark of *Stryphnodendron adstringens* (“barbatimão”), another native species of the Brazilian Caatinga, also increased the viability of HaCaT (Pellenz et al., 2018). In the scratch assay, MFSOL showed a proliferative and migratory effect on keratinocytes during wound healing. In this sense, a study with leaves of *Aloe vera*, *Aloe ferox* and *Aloe marlothii* presented rapid coverage rates in the scratch assay on HaCaT keratinocytes and low cellular cytotoxicity (Fox et al., 2017).

To evaluate the isolate effect on cell migration, mitomycin C, an antimitotic agent was used to interfere with cell duplication in the scratch assay. Thus, MFSOL (25 and 50 μg/ml) stimulated not only proliferation but also the migration of HaCaT. A similar result was observed by the methanol fraction of the extract of *Centella asiatica* (100 μg/ml) that increased the cell migration rate, besides assisting in the remodeling during cicatrization of excisional wounds in rabbits (Azis et al., 2017). Epidermal cell turnover is fundamental for rapid wound healing, in which keratinocytes are stimulated to proliferate by regulating growth factors and intercellular contact (J. H. Choi et al., 2017).

One of the parameters of success in healing is reepithelialization. When keratinocytes fail to recover the epidermal barrier, new openings may occur, especially in chronic lesions (Rousselle et al., 2018). As a pharmacological model, the induction of superficial burn in mice was chosen to evaluate the effect of a topical cream (CrMFSOL) containing MFSOL at concentrations of 0.5% and 1.0% for 14 days. The faster evolution of the contraction area on burn wounds, evidenced by treatment with CrMFSOL after 3, 5, 7 and 9 days, did reduce the risk of infections and accelerated the other stages of healing, such as proliferation and remodeling. Its effect was similar to the control group (Silver Sulfadiazine), the standard treatment for the prevention of infections and the promotion of cicatrization in burns (Moser, Pereima, & Pereima, 2013). A methanol extract of *Carissa spinarum* (1.0%) used in the treatment of partial burns in rats also showed a more pronounced effect on the contraction of the lesion in the first days evaluated, at the same concentration that MFSOL (Sanwal & Chaudhary, 2011).

The topic treatment with CrMFSOL stimulated the early formation of a new epithelium and a remodeling of connective tissue, with a higher collagen deposition after 7 days. Similarly, a phytomodulatory hydrogel with hemicelluloses extracted from seeds of *Caesalpinia pulcherrima* (Fabaceae) and mixed with phytomodulatory proteins obtained from the latex of *Calotropis procera*, enhanced the healing effects by high collagen deposition and modulating some aspects of the inflammatory phase (Vasconcelos et al., 2018). Corroborating with this data, it was discovered that the methanol fraction of quixaba leaves (MFSOL), contains as a main component, the NMP, a derivative from L-proline (amino acid precursor of collagen), that showed cicatrizant effect in the topical treatment of induced excisional lesions in mice, through modulation of the inflammatory response, and increase of the antioxidant response and stimulation of the production of collagen fibers (Aquino et al., 2019). Also, a study with the ethanol extract of the leaves of *Passiflora edulis* reproduced an acceleration the repair, to reduce the number of inflammatory cells on the 7th day of treatment and to increase the number of fibroblasts and the deposition and organization of the fibers of collagen in 14 days of repair of burns in mice (Barros, Santos, Coelho, Reis, & Bezerra, 2016).

The progression of thermal injury damages adjacent capillaries causing ischemia, which activates adhesion of polymorphonuclear cells, such as neutrophils, which release inflammatory mediators as well as the production of reactive oxygen species (ROS) (Parihar, Parihar, Milner, & Bhat, 2008). While TNF-α release induces the fundamental inflammatory response in damaged tissue, the high level of IL-1β in mouse burns was correlated to the increased activity of epidermal keratinocytes. These cells work to restore the epidermal barrier function when stimulated by nanofibrous of peptides hydrogels, (Loo et al., 2014). A biomembrane containing latex proteins from *Calotropis procera* increased levels of TNF-α and IL1-β in the model of excisional wounds in mice and promoted an improvement in wound healing (Ramos et al., 2016). Also, CrMFSOL 0.5% treatment provided an increase in IL-1β levels, but it did not influence on TNF-α release on burn injuries. Moreover, CrMFSOL 0.5% reduced MPO activity, after 3 days of treatment, which may have influenced the transition from acute to chronic inflammatory infiltrate after 7 days earlier than the other treatment with Silver Sulfadiazine.

Besides, IL-1β has been reported to stimulate VEGF expression during inflammation, which consequently stimulates migration and proliferation of endothelial cells by the promotion of angiogenesis. Thus, the healing effect of a polysaccharide extracted from *Sanguisorba officinalis* to induce epithelization and angiogenesis was related to the increase of VEGF and IL-1β in mouse burn model (Zhang, Chen, & Cen, 2018). Similarly, CrMFSOL 0.5% promoted a greater release of VEGF after 14 days of treatment on burn injuries, which may have contributed to a better epithelization and reorganization of the connective tissue.

As an anti-inflammatory factor, IL-10 is also produced by epidermal keratinocytes and is important in the suppression of ROS, Nitric Oxide (NO) and the control of proinflammatory cytokines secreted by macrophages (Tharuka et al., 2018). Briefly, the CrMFSOL 0.5% provided an increase in IL-10 levels, MFSOL was able to modulate the inflammatory response and contribute to recovering of tissue damage promoted by burn injury.

## 5. CONCLUSION

The methanol fraction extracted from the leaves of *Sideroxylon obtusifolium* demonstrated wound healing potential on stimulating proliferation and migration of human keratinocytes. Also, MFSOL was able to modulate the inflammatory response and improve the tissue repairment of superficial burns in mice. Thus, it is believed that the anti-inflammatory and wound healing potential of MFSOL may provide support for further studies that enable the development of an herbal product indicated to the treatment of wounds and burns.

## Acknowledgments

This study was financially supported by CNPq (Conselho Nacional de Desenvolvimento Científico e Tecnológico). We acknowledge the support from Drug Research and Development Center (NPDM) for the pharmacological tests.

